# Unleashing meiotic crossovers in crops

**DOI:** 10.1101/343509

**Authors:** Delphine Mieulet, Gregoire Aubert, Cecile Bres, Anthony Klein, Gaëtan Droc, Emilie Vieille, Celine Rond-Coissieux, Myriam Sanchez, Marion Dalmais, Jean-Philippe Mauxion, Christophe Rothan, Emmanuel Guiderdoni, Raphael Mercier

**Affiliations:** CIRAD, UMR AGAP, 34398 Montpellier Cedex 5, France; Univ Montpellier, CIRAD, INRA Montpellier SupAgro, Montpellier France; Agroécologie, AgroSup Dijon, INRA, Univ. Bourgogne Franche-Comté, F-21000 Dijon, France; UMR 1332 BFP, INRA, Univ. Bordeaux, F-33140 Villenave d’Ornon, France; Institue of Plant Sciences, Paris Saclay IPS2, CNRS, INRA, Université Paris-Sud, Université Evry, Université Paris-Saclay, 91405 Orsay, France; Institut Jean-Pierre Bourgin, UMR1318 INRA-AgroParisTech, Université Paris-Saclay, RD10, 78000 Versailles, France.

## Abstract

Improved plant varieties are hugely significant in our attempts to face the challenges of a growing human population and limited planet resources. Plant breeding relies on meiotic crossovers to combine favorable alleles into elite varieties^1^. However, meiotic crossovers are relatively rare, typically one to three per chromosome^2^, limiting the efficiency of the breeding process and related activities such as genetic mapping. Several genes that limit meiotic recombination were identified in the model species Arabidopsis^2^. Mutation of these genes in Arabidopsis induces a large increase in crossover frequency. However, it remained to be demonstrated whether crossovers could also be increased in crop species hybrids. Here, we explored the effects of mutating the orthologs of *FANCM*^3^, *RECQ4*^4^ *or FIGL1*^5^ on recombination in three distant crop species, rice (*Oryza sativa*), pea (*Pisum sativum*) and tomato (*Solanum lycopersium*). We found that the single *recq4* mutation increases crossovers ~three-fold in these crops, suggesting that manipulating *RECQ4* may be a universal tool for increasing recombination in plants. Enhanced recombination could be used in combination with other state-of-the-art technologies such as genomic selection, genome editing or speed breeding^6^ to enhance the pace and efficiency of plant improvement.

Meiotic crossovers shuffle chromosomes to produce unique combinations of alleles that are transmitted to offspring. Meiotic crossovers are thus at the heart of plant breeding and any related genetic analysis such as quantitative trait loci (QTLs) detection or gene mapping. However, crossovers are relatively rare events, which is intriguing since their molecular precursors (*i.e.* DNA double stranded breaks and inter-homologue joint molecules) largely outnumber the final crossover number. Indeed, It was recently shown that active mechanisms limit the formation of meiotic crossovers in Arabidopsis ^2–5,7–9^. Forward genetic screens identified three anti-crossover pathways that rely on the activity of FANCM, RECQ4 and FIGL1, respectively. RECQ4 appears to be the most important anti-crossover factor, as the mutation of the corresponding genes (*RECQ4A* and *RECQ4B*) led to an almost four-fold increase in recombination in Arabidopsis hybrids^2,10^. RECQ4 is a DNA helicase homologue of mammalian BLOOM and yeast Sgs1^11,12^. *FANCM*, which encodes another conserved DNA helicase, is also an important anti-crossover factor in Arabidopsis. Mutation of this gene also leads to a large increase in recombination, but only in pure lines (~3-fold) with a very limited effect in hybrids ^2,3,5,13^. FANCM was also shown to limit crossovers in a *Brassica rapa* pure line ^14^. The third pathway depends on the AAA-ATPase FIGL1. Mutation in *FIGL1* alone leads to a relatively modest increase in recombination (+25% in Arabidopsis hybrids), but when combined with *recq4*^2^ it leads to an almost eight-fold increase. Mutation in *FIGL1* leads to full sterility in rice^15^, raising doubts about the pertinence of manipulating this gene in crop species.

Here we tested the effect of *recq4*, *fancm* and *figl1* mutations on recombination in three crop species. We chose rice (*Oryza sativa*), the cultivated pea (*Pisum sativum*) and tomato (*Solanum lycopersium*) for their economic importance and because they represent distant clades. Indeed they are members of the three major clades of flowering plants, monocots, eudicots rosids and eudicots asterids, respectively ^16^. Rice is the staple of more than half of mankind and as such is the number one cereal consumed. It belongs to the *Poaceae* family that also contains maize, wheat and barley ^16^. Pea, in addition to be the genetic model used by Mendel, is the second most cultivated pulse crop in the world (http://faostat.fao.org/) and belongs to the *Fabaceae* family that contains many crop species such as chickpea, beans and lentil. Tomato, the second most cultivated fresh-market vegetable crop, is one of the most important nutrient-dense superfoods and belongs to the *Solanaceae* family, which includes potatoes, eggplant and peppers.

We first explored the conservation and copy number of *RECQ4*, *FANCM* and *FIGL1* in flowering plants (Figures S1-3, Dataset S1). For *FANCM* and *FIGL1*, a single homolog of each gene was identified in most species including pea, tomato and rice. Several copies of *FANCM*^17^ and *FIGL1* were found only in very recent polyploids (e.g wheat). Several copies of *RECQ4*, on the other hand, appear to have been retained from earlier whole genome duplications in several clades, leading to the presence of two or more copies in several species (e.g. Arabidopsis^4,11^, cabbage, lettuce, soybean, sunflower).

We then assessed the role of *OsRECQ4* (Os04g35420) and *OsFANCM* (Os11g07870) in meiotic recombination in rice. We screened mutant collections of two different cultivars, Nipponbare ^18,19^ and Dongjin ^19^, that are both from the *japonica* temperate sub group. Comparison of 25X Illumina sequencing of Dongjin and the Nipponbare reference genome, showed a divergence of one single nucleotide polymorphism (SNP) per ~11kb (M&M). We identified one insertion mutant in each cultivar for both genes (Figures 1A and S4). As mentioned above, mutation of *FIGL1* was recently shown to cause sterility in rice and was thus not further studied here^15^. We produced Dongjin/Nipponbare F1 hybrids mutant for both *OsRECQ4* alleles, or for both *OsFANCM* alleles and wild type siblings (M&M and Figure S5). Hybrid fertility was not affected by either the bi-allelic *Osfancm* or the *Osrecq4* mutation (Figure 1B). No defects in meiosis progression were observed during male meiosis in *Osrecq4* hybrids (Figure S6), which is consistent with normal fertility. F1 plants were self-fertilized to generate F2 populations that were genotyped for an average of 19 SNP markers per chromosome (on the 12 chromosomes for the *RECQ4* populations and five chromosomes for the *FANCM* populations). We analyzed 149 *Osrecq4 -/-* F2 plants, 108 *Osfancm -/-* F2 plants and a total of 262 wild types (Dataset S2). In *Osrecq4 -/-*, we observed an increase in the genetic size of all 12 chromosomes leading to a 3.2-fold increase in the total genetic map compared to wild type (total size ± 95% confidence interval: 5700 ± 231 cM vs 1759 ± 58 cM;) (Figures 2 and 3). This shows that RECQ4 is a major meiotic anti-crossover factor in hybrid rice. In *Osfancm -/-*, recombination was increased by 2.3-fold (cumulated genetic map size of the five chromosomes analyzed: 1649 ± 122 cM vs 724 ± 69 cM in wild type). This is remarkable, as no increase in recombination was observed in Arabidopsis *fancm* hybrids^2,10^ (Figures 2 and 3). Crossover distribution along the chromosomes (Figure 4) showed that in both *Osfancm* and *Osrecq4*, enhancement of recombination occurs along chromosome arms but not in the peri-centromeric regions, suggesting that other factors limit crossovers in these regions, as previously proposed for Arabidopsis. In addition to peri-centromeres, another region on the right arm of chromosome 11 was relatively supressed for crossovers in wild type and *Osrecq4* (Figure 4). Interestingly, this region is associated with a cluster of resistance genes^20^ and diverges significantly between the parental genomes. The same observation was made in Arabidopsis ^2,10^. This suggests that regions with high levels of polymorphism are less prone to the extra crossovers that arise in *recq4* mutants.

**Figure 1A.**
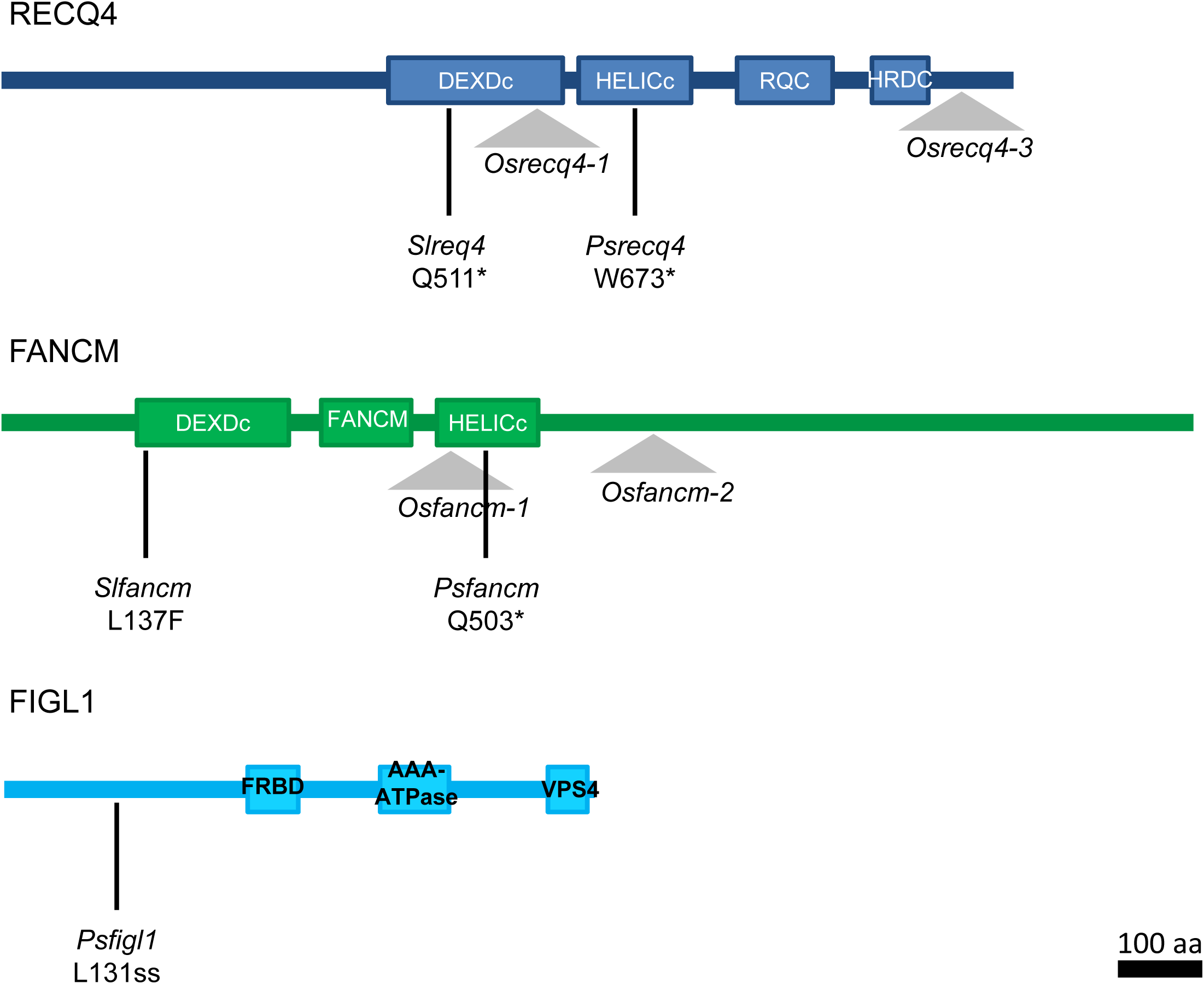
Graphical representation of the RECQ4, FANCM and FIGL1 proteins and positions of the mutations described in this study for rice (Os for *Oryza sativa*), tomato (Sl for *Solanum lycopersicum*) and pea (Ps for *Pisum sativum*). T-DNA insertions are indicated with a triangle and EMS point mutations with a black vertical line. Conserved Protein domains are represented by rectangles. AAA-ATPase: ATPase Associated with diverse cellular Activities; DEXDc: DEAD-like helicase domain; FANCM: Fanconi anemia complementation group M; FRBD: FIDGETIN-RAD51-Binding-Domain; HELICc: Helicase superfamily C-terminal domain; HRDC: Homologous region RNase D C-terminal; RQC: RecQ C-terminal; VPS4: Vacuolar Protein Sorting 4.

**Figure 1B.**
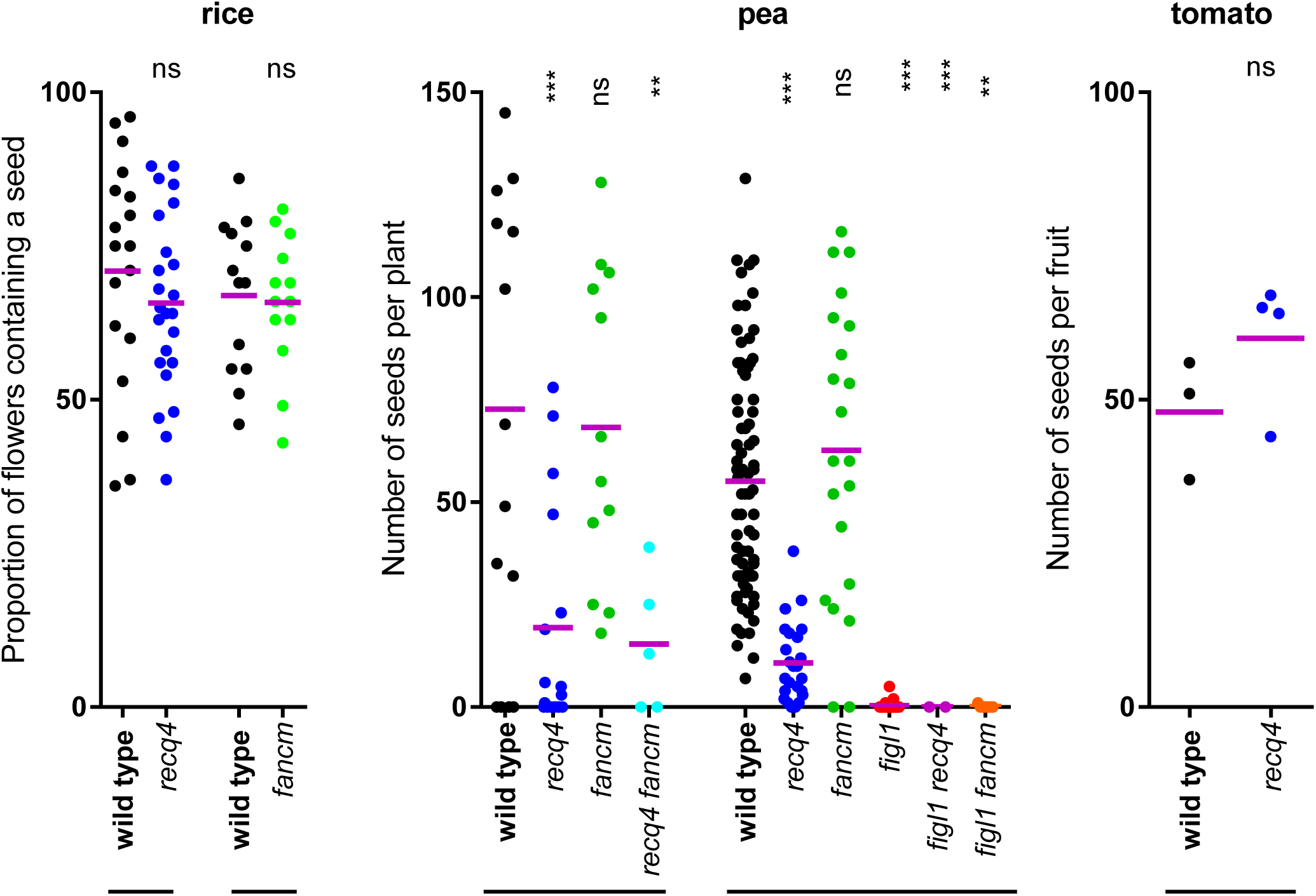
Fertility Analysis of the *recq4*, *fancm* and *figl1* mutants in rice, pea and tomato For rice, each dot represents the fertility of an individual plant measured as the proportion of flowers giving rise to a seed (n>150 flowers/plant). For pea each dot represents the fertility of an individual plant measured as the total number of seeds per plant. For tomato each dot represents the fertility of an individual plant measured as the number of seeds per fruit (n=3 fruits per plant). The bar under the graph indicates that the plants are siblings. The purple bars represents the mean. Anova with Sidaks’s multiple comparison correction: *** p<0.001; ** p<0.01; not significant (ns) p>0,05.

**Figure 2.**
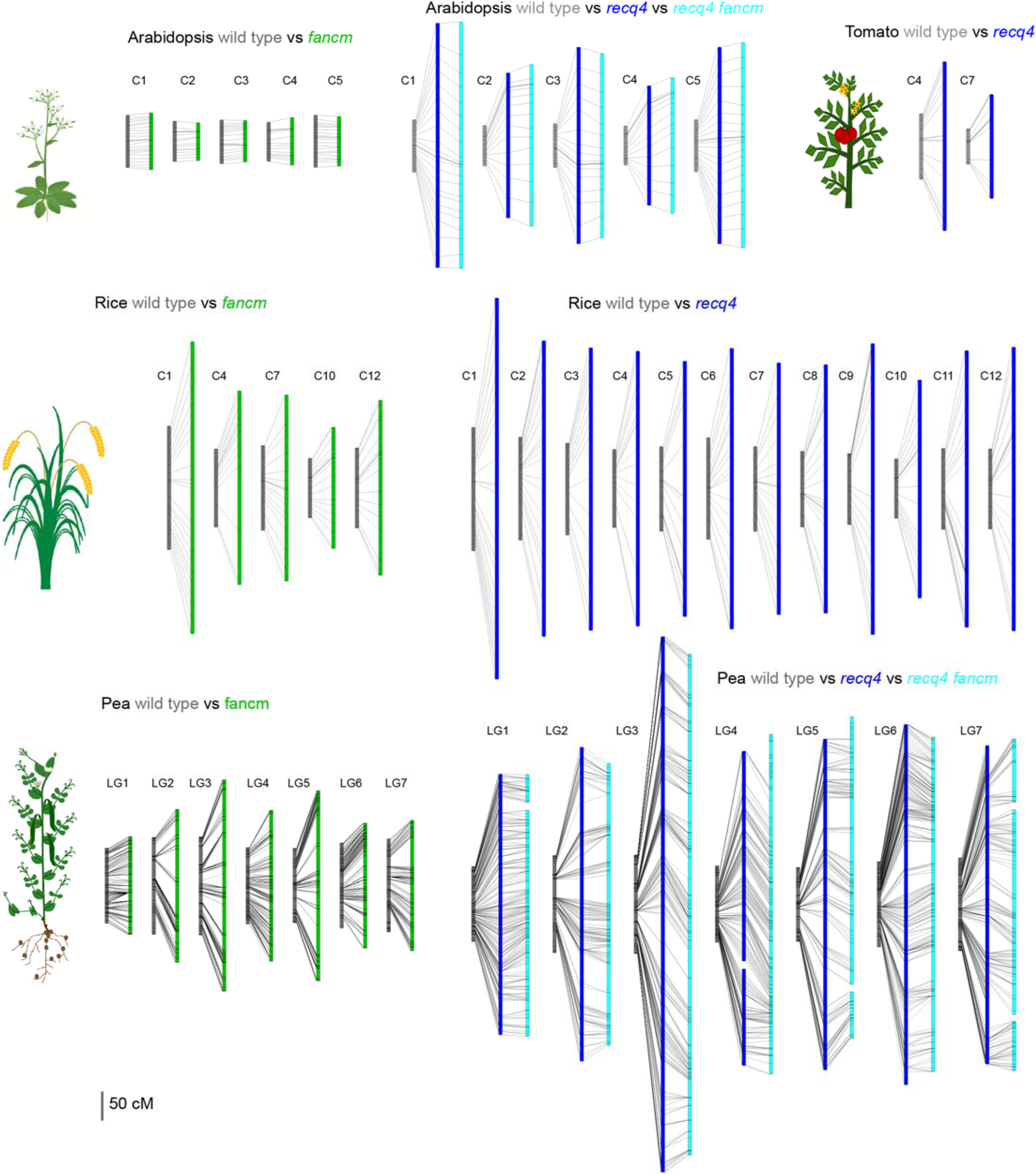
Genetic maps in *fancm* and *recq4* mutants compared with wild type for Arabidopsis, rice, pea and tomato. C=chromosome. LG=Linkage group. Each black line represents an informative genetic marker. Data can be found in Tables S3, S5 and S7. Data for Arabidopsis are from Fernandes et al ^2^

**Figure 3.**
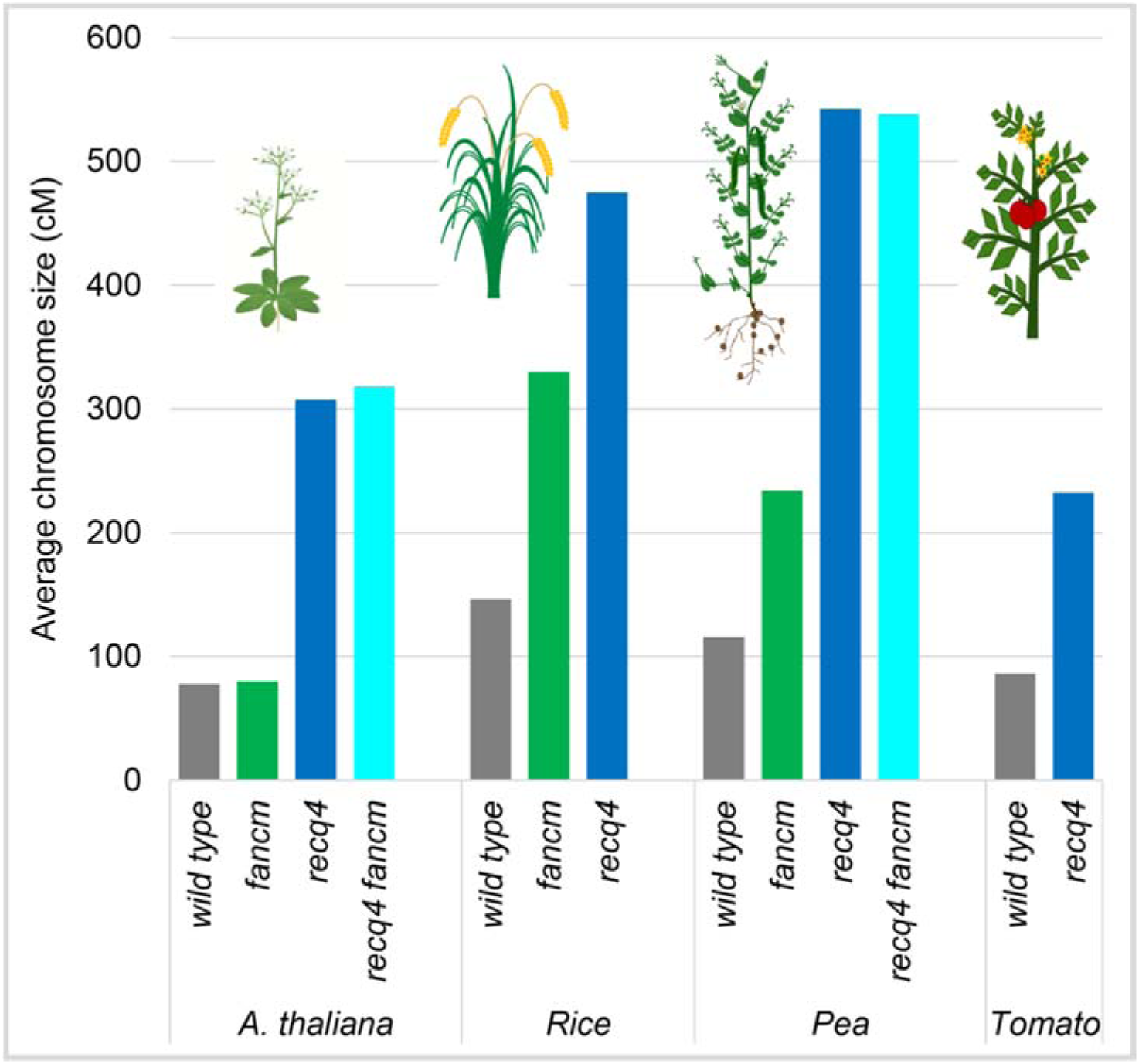
Average chromosome size in wild type, *fancm* and *recq4* mutant plants for Arabidopsis, rice, pea and tomato.

**Figure 4.**
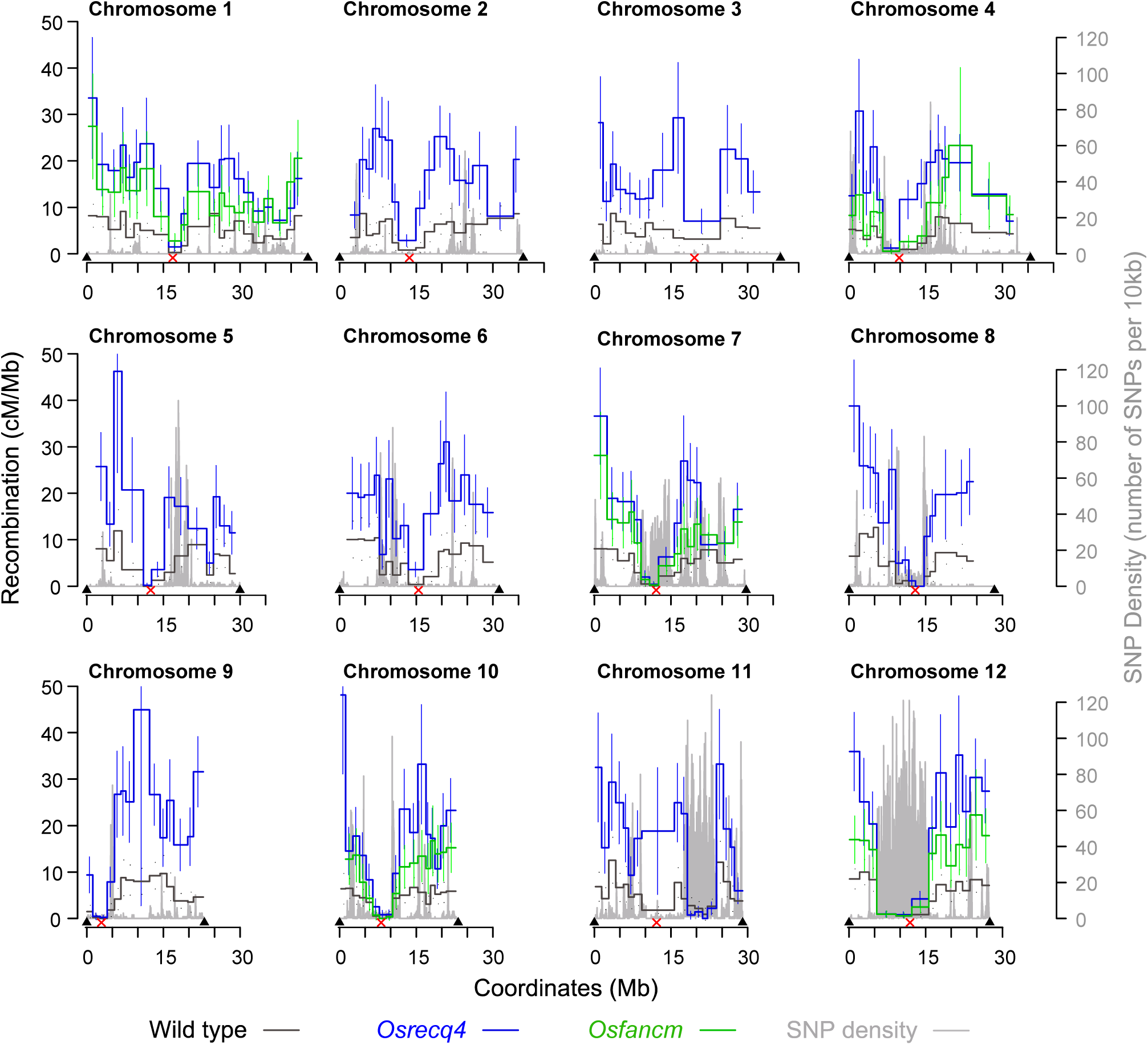
Distribution of COs along the 12 rice chromosomes in Osrecq4l (blue), Osfancm (green) and wild type (grey) plants. The recombination frequency (cM/Mb) in each interval was plotted along the 12 rice chromosomes. The density of SNP polymorphisms between Dongjin and Nipponbare strains is shown in grey. Red crosses represent the centromere positions; the arrows represent the telomere positions.

Next, we extended our analysis to the pea *Pisum sativum* by screening an EMS-induced mutant population^21^ (cultivar Cameor). We identified a STOP-codon mutation in *P*s*FANCM*, *PsRECQ4* and a splicing site alteration in *PsFIGL1* (*fancm-Q503\**, *recq4-W673*, figl1-L131ss;* Figure 1A). To measure the effect of these mutations alone or in combination, we produced two independent populations. The first population segregated the *Psrecq4* and *P*s*fancm* mutations and genetic polymorphisms from a different cultivar (Kayanne) (M&M and Figure S7). The second population was purely Cameor and segregated the three mutations (Figure S8). Fertility was quite variable from plant to plant, presumably because of the segregation of additional EMS mutations. In both populations, the fertility of F2 *Psfancm* mutants was indistinguishable from that of wild type. However, all the F2 plants that were homozygous for the *Psfigl1* mutation were sterile (Figure 1B) and *Psrecq4* mutants produced four times less seed than wild type. This suggests that PsFIGL1 is essential for meiosis and fertility in pea, as previously shown in rice^15^, and that PsRECQ4 may also be required for full fertility (Figure 1B). However, we cannot rule out the possibility that this reduced fertility in *Psrecq4* and *Psfigl1* was caused by additional linked EMS mutations. Seeds could be obtained in sufficient numbers for *P*s*recq4*, *P*s*fancm*, *P*s*recq4 P*s*fancm* and wild type siblings (Figure S7). For each of these genotypes, ~50 F3 plants were genotyped for 5097 SNPs between the cultivars Cameor and Kayanne (Dataset S4) to measure genome wide recombination. Note that because certain regions were fixed in the F2s, only ~80% of the genome was segregating for polymorphic markers in the four genotypes and was thus analyzed to compare recombination levels (810 cM of the 1018 cM of the total wild type map). For *P*s*fancm*, we observed a global twofold increase in recombination (1639 ± 204 cM vs 810 ± 78 cM), similar to that observed in rice, but in contrast with the absence of effect in Arabidopsis hybrids. In *Psrecq4*, recombination increased even further with 4.7 times more crossovers observed compared to wild type (3798 ± 296 cM vs 810 ± 78 cM) (Figures 2 and 3). Thus RECQ4 is a major anti-crossover factor in Pea. *Psrecq4* and *Psfancm* double mutants did not show a further increase in recombination compared to *Psrecq4* alone (3767 ± 288 cM vs 3798 ± 296 cM). This suggests that in Pea either PsRECQ4 and PsFANCM act in the same anti-crossover pathway, which would be intriguing as these two helicases appear to act in parallel in Arabidopsis, or that some upper limit has been reached (e.g. the use of all eligible crossover precursors).

Finally, we looked for mutations in *FANCM* and *RECQ4* in a tomato EMS-induced mutant population (Cultivar Micro-Tom) (Figure 1A). We identified a STOP codon in *SlRECQ4* (*recq4-Q511\**) and crossed the corresponding line to wild type cultivar M82 (M&M and Figure S9). Wild type and *Slrecq4* F2 plants had similar fertility (Figure 1B). We focused our analysis on chromosome 4 and 7 and observed a 2.7-fold increase in recombination in the mutant compared to the wild type (cumulative map 173 ± 22 cM vs 464 ± 52 cM) (Figures 2 and 3). This shows that RECQ4 is also a major factor limiting meiotic recombination in tomato. We also identified missense mutations in tomato *FANCM* in a conserved amino acid (L137F). Following a similar approach as described above for *recq4*, we did not detect an increase in meiotic recombination in hybrids homozygous for this mutation (data not shown). However, further work is needed to understand whether disruption of *FANCM* has no effect in this context, as observed for Arabidopsis hybrids, or if the L137F mutation does not fully disrupt FANCM activity.

## Discussion

Here we explored the potential for *fancm* and *recq4* mutation to increase recombination in crops. In Arabidopsis, the *fancm* mutation leads to a threefold increase in recombination in a pure line but has almost no effect in hybrids (Col/Ler)^5,13,22^. However, we showed here that mutating *FANCM* results in a ~twofold increase in recombination in hybrid rice (Dongjin/Nipponbare) and hybrid pea (Cameor/Kayanne). This difference could be due to variation in the recombination machinery in these species or be associated with the level of polymorphisms in these hybrids. Indeed, the SNP density is ~1/200pb in the Col-Ler Arabidopsis hybrid^23^, but is much lower in the rice Dongjin/Nipponbare (1/11kb) and Cameor/Kayanne pea hybrids (~1/10kb and ~1/5kb, respectively) and, by definition, virtually null in the Arabidopsis pure line. This would mean that the *fancm* mutation only increases recombination if the polymorphism rate is below a certain threshold, somewhere between 1/200 and 1/5000 SNPs per kb. It would be interesting to explore the *fancm* effect in more distant hybrids (e.g. *Japonica-Indica* rice) or in different species, to test this hypothesis.

We showed that the *recq4* mutation alone can massively increase recombination in rice, pea and tomato hybrids, a result similar to that observed in Arabidopsis^2^. This suggests that mutation in *RECQ4* orthologs may be a universal approach for enhancing recombination rates in crop species. These increases in crossover frequency are much higher than any previously observed natural or environmentally-induced variation in recombination (e.g. temperature which typically modifies recombination by 10-30% ^24–27^). Increased recombination is predicted to improve the response to selection in the short, medium, and long term^28^. Thus higher recombination rates could be used to enhance genetic gain in breeding schemes. Further, increased recombination would also enhance the power of pre-breeding activities such as genetic map construction, QTL detection, and positional cloning. However, the *recq4* mutation does not homogeneously increase recombination along the genome (Figure 3 and ^2^). First, the peri-centromeric regions, that are reluctant to crossover in wild type, still fail to recombine in the mutants, suggesting that additional unknown mechanisms prevent crossovers close to centromeres ^29^. Future studies should prioritize the identification of these mechanisms and methods to increase crossover in proximal regions as these regions represent a large part of the genome in important crops such as wheat^30^. Second, the increase in recombination tends to be lower in more divergent regions of the genome. Strikingly, the regions of highest sequence divergence showed a limited increase in recombination compared to the rest of the genome (Figure 4 and ^2^). This suggests that the extra crossovers arising in the *recq4* or *fancm* mutants tend to be prevented by sequence divergence. This predicts that mutating *recq4* could be ineffective for promoting recombination between distant genomes, such as in interspecific crosses, but this remains to be tested. The same appears to be true for all anti-crossover genes identified to date^7,10^. Further studies are required to understand how sequence divergence drives genetic recombination.

In all species examined so far, mutation in *RECQ4* resulted in the most significant increases in crossover numbers. However in Arabidopsis, further increases were obtained by combining the *recq4* mutation with either a mutation in the *FIGL1* gene, or with overexpression of HEI10 ^2,10^. While *figl1* only mildly affects fertility in Arabidopsis, it leads to sterility in rice ^15^ and pea, precluding the use of *figl1* to manipulate recombination in those species. Both *figl1* mutation and HEI10 overexpression remain to be tested in other species.

Here we used classic mutagenesis to disrupt *FANCM* and *RECQ4* and crosses to introduce this mutation into the hybrid context. However, the development of very effective targeted mutagenesis techniques based on CRISPR-cas9 now offers the possibility to disrupt these genes directly in the F1 hybrids ^31^ and thus rapidly obtain hyper-recombined populations and enhance the efficiency of crop breeding.

## Materials and methods

### Phylogeny

Sequences from RECQ4, FANCM and FIGL1 proteins were retrieved from the PLAZA V4 dicots and monocots databases^32^ https://bioinformatics.psb.ugent.be/plaza/ using BLASTP (RECQ4: ORTH004M000654 and ORTH004D00423; FANCM ORTHO04D004865 and ORTHO04M004526) and species by species using BLASTP on the nr database at NCBI (https://blast.ncbi.nlm.nih.gov/Blast.cgi?PAGE=Proteins), Phytosome12 (https://phytozome.jgi.doe.gov/pz/portal.html), the Pea RNA-Seq gene atlas (http://bios.dijon.inra.fr/FATAL/cgi/pscam.cgi)^21^, the Sol Genomics Network https://solgenomics.net/ ^33^ and the IWGSC RefSeq Annotations. For each candidate gene, if several protein isoforms/predictions were present in the databases, the isoform/prediction with the higher similarity to the corresponding protein in other species was retained for further analysis (Dataset S1). The phylogenetic analysis was performed on the Phylogeny.fr platform^34^ and included the following steps: 1) Sequences were aligned with MUSCLE (v3.8.31) configured for highest accuracy; 2) Positions with gaps were removed from the alignment; 3) The phylogenetic tree was reconstructed using the maximum likelihood method implemented in the PhyML program (v3.1/3.0 aLRT). The default substitution model was selected assuming an estimated proportion of invariant sites (of 0.118) and four gamma-distributed rate categories to account for rate heterogeneity across sites. The gamma shape parameter was estimated directly from the data (gamma=0.929). Reliability of internal branches was assessed using the aLRT test (SH-Like). Graphical representation and editing of the phylogenetic tree were performed with TreeDyn (v198.3) and adobe illustrator.

### Rice

Illumina Paired-end reads from Dongjin were aligned to the Nipponbare reference genome (MSU7) using the software BWA (release 0.7.10). PCR artifacts were removed by Picard tools MarkDuplicates (https://broadinstitute.github.io/picard/). SNP and INDEL identification were performed with GATK HaplotypeCaller (release 3.4-0-g7e26428) with default parameter. Raw variants were filtered according to GATK recommendations (https://software.broadinstitute.org/gatk/best-practices/). In resulting VCF file (Variant Call Format) we retained only variants that have passed all filters (PASS quality) and we selected homozygous SNPs (both alleles are different from those of Nipponbare reference). 33540 SNPs were retained for a total genome size of 373 Mb, corresponding to 1 SNP per 11 Kb between Dongjin and Nipponbare cultivars.

The following mutations were used in this study: *Osfancm-1* (AQSG07), *Osfancm-2* (A46543), *Osrecq4l-1* (3A-03503) and *Osrecq4l-3* (AUFG12) (Figure 1A and Figure S4). *Osfancm-1* and *Osrecq4l-3* are in the Nipponbare cultivar from the Oryza Tag Line insertion line library ^18,35^. *Osfancm-2* and *Osrecq4l-1* are in the Dongjin cultivar from the POSTECH Rice Insertion Database ^36^. For each allele, the position of the T-DNA in the rice genome was confirmed with Sanger sequencing (Figure S4). Plants were grown under containment greenhouse conditions (28°C / 24°C day/night cycle, 60% humidity) with natural light boosted by artificial sodium lights (light intensity of 700 µmoles/m2/s). The crossing scheme is summarized in figure S5. Heterozygous plants for the mutations were identified using PCR. Primers were designed using the “Genotyping Primer Designer” tool of OryGenesDB (http://www.orygenesdb.cirad.fr)^35,37^. Genotyping primers and expected PCR product sizes are listed in Table S1. We crossed the heterozygous lines *Osfancm-2+/-* with *Osfancm-1+/-* and *Osrecq4-1+/-* with *Osrecq4-3+/-*. Dongjin lines were used as female and Nipponbare lines as male (Figure S5). Crosses were carried out through manual castration of florets and pollination, followed by bagging to avoid pollen contamination. F1 hybrid plants were genotyped twice to select *Osfancm*-/-, *Osrecq4*-/- and their respective wild type siblings (Figure S5). F1 sibling plants of the desired genotypes were used for fertility measurements, cytological analyses and selfed to produce the F2 populations. Male meiotic chromosome spreads were performed as previously described^38^. For SNPs genotyping of the F2s, DNA was extracted from 500mg of fresh leaves and adjusted to 10ng/µL. Single nucleotide polymorphism genotyping was performed using Kompetitive Allele Specific PCR (KASP) following the LGC group recommendations for the use of KASP technology on Biomark Fluidigm with a set of 241 robust KASP markers spread over the physical map (~every 1.5 Mb). Genotyping data were analyzed with Fluidigm software (Fluidigm SNP Genotyping Analysis 4.3.2) with manual error corrections. The raw genotyping dataset is shown in Dataset S2. Recombination analysis was performed with MapDisto 2.0 b105 ^39^. Linkage groups were determined for the wild type F2 population (LOD1, RFmax: 0,5), and fit perfectly with the physical marker order. Genotyping errors were filtered using the iterative error removal function (iterations = 5, start threshold = 0.001, increase = 0.001). Recombination (cM ± SEM) was calculated using classical fraction estimation and the Haldane mapping function. The obtained recombination frequencies per interval and corresponding genomic data are shown in Dataset S3. Graphical representations were generated with R 3.3.2 (Figure 4).

### Pea

Mutations in *PsRECQ4*, *PsFANCM* and *PsFIGL1* were identified using TILLING (Targeting Induced Local Lesions IN Genome) in the cultivar Cameor, and combined by crosses. In the *Psfancm* mutant there is a C to T transition at position 1507 from the A of the start codon of the coding sequence, leading to a nonsense mutation (Q503*). In *Psrecq4* there is a G to A transition at the position 2019 from the A of the start codon of the coding sequence, leading to a nonsense mutation (W673*).

The *PsFigl1* mutation is a G to A transition at position 3740 from the ATG on the genomic sequence, modifying the splice junction before the 3rd exon. Two independent populations were produced (Figure S7 and S8).

In the first population, one plant *PsRECQ4+/- PsFANCM+/-* was crossed to the wild type cultivar Kayanne (Figure S7). One F1 plant was selfed to produce 180 F2 plants, among which single mutants, double mutants and wild type were identified by genotyping. Five *Psfancm*, five *Psrecq4*, three *Psfancm Psrecq4* and five wild type F2 plants were selfed to produce the F3 populations (~50 plants per genotype).

In the second population, two Cameor *PsRECQ4+/- PsFANCM+/- PsFIGL1+/-* were selfed to produce 160 F2 plants (Figure S8). Twenty-one *Psfancm* mutants, 24 *Psrecq4,* 24 *Psfigl1,* 2 *Psfigl1Psrecq* double mutants and 7 *Psfigl1Psfancm* double mutants were identified by genotyping. Fertility was analyzed for the two F2 populations (Figure 1 B).

F2 and F3 plants of the Cameor/Kayanne hybrid population were genotyped for 5097 markers polymorphic between Kayanne and Cameor using the GenoPea 13.2K SNP Array^40^ (Dataset S4). Markers that were homozygous in F2 plants were scored as missing data in its F3 progeny. Very rare dubious singletons were manually edited into missing data. Recombination analysis was carried out with MapDisto 2.0 b104 ^39^, using the linkage groups defined in ^40^ with some manual corrections that minimized the number of crossovers. The F2 and F3 wild type maps were not significantly different from each other and were combined to gain detection power. Recombination (cM ± SEM) was calculated using classical fraction estimation and the Haldane mapping function. The obtained recombination frequencies per interval and corresponding genomic data are shown in Dataset S5. Complete maps are shown in figure 2. Only the genetic space for which data were obtained in the four genotypes (~80% of the total map) is shown in figure 3 (common map in Dataset S5).

### Tomato

Q511>STOP *RECQ4* and L137F *FANCM* mutations were isolated using TILLING in a tomato EMS mutant collection in the cultivar Micro-Tom ^41,42^. Genetic mapping was carried out in F3 populations from a cross between a *reccq4-Q511\** homozygous mutant and the processing variety M82 (Figure S9). A 96 F2 population from a F1 hybrid was genotyped for the *recq4* mutation using a set of 30 markers on chromosomes 4 and 7 that are polymorphic between Micro-Tom and M82^43,44^ (Dataset S6). A total of 16 F2 plants were selected for their maximal heterozygosity for chromosome 4 or chromosome 7 and for being either *RECQ4*+/+ or *recq4*-/- (Figure S9). Forty F3 progenies were generated by selfing from each of these F2 plants. The 640 F2 plants were genotyped for SNP markers on chromosome 4 or 7. The plants were grown and DNA extracted as described in ^43,45^. Genotyping was performed by KASP^TM^ Assay^46^. Markers that were homozygous in F2 plants were scored as missing data in its F3 progeny. Recombination analyses were performed with MapDisto 2.0 b104^47^. Genotyping errors were filtered using the iterative error removal function (iterations = 1, start threshold = 0.001). Recombination (cM ± SEM) was calculated using classical fraction estimation and the Haldane mapping function. The obtained recombination frequencies per interval are shown in Dataset S7.

## Acknowledgment

We thank Judith Burstin, Mathilde Causse, Brigitte Courtois and Christine Mézard for fruitful discussions. We thank Pierre Sourdille and Fatiha Benyahya for sharing wheat sequences before publication. We thank Christine Le Signor and Marie-Christine Le Paslier for offering their expertise and advices. The pea and tomato work was funded by the HyperRec grants from INRA Transfert. The Institute Jean-Pierre Bourgin benefits from the support of the LabEx Saclay Plant Sciences-SPS (ANR-10-LABX-0040-SPS). This work was partly funded by the “Investissements d’Avenir”, France Génomique project IRIGIN (INTERNATIONAL RICE GENOME INITIATIVE).

---

**Figure S1.**
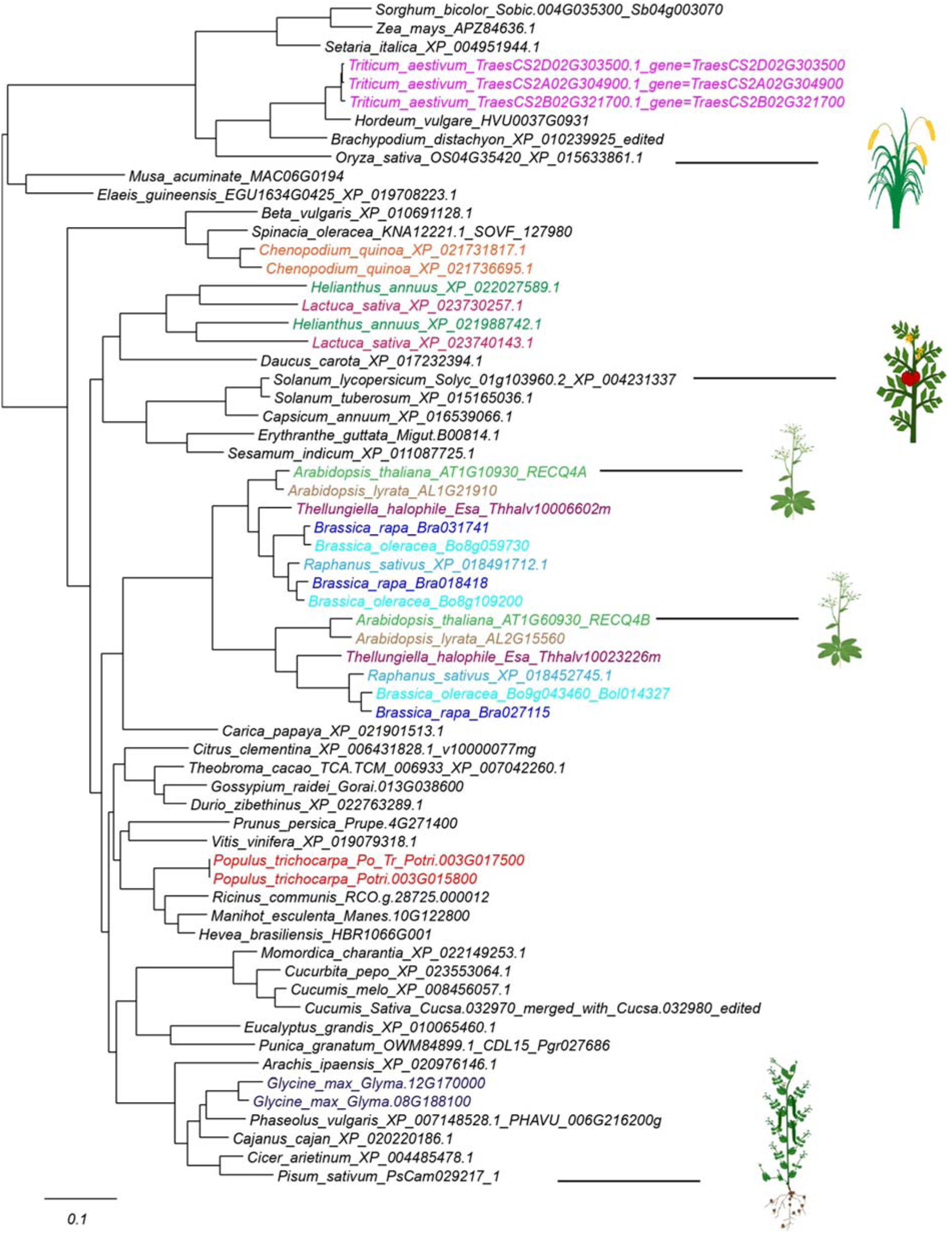
Phylogenetic tree of plant RECQ4 proteins. Genes present in several copies in a given species have been colored. Proteins sequences and accession numbers can be found in dataset S1

**Figure S2.**
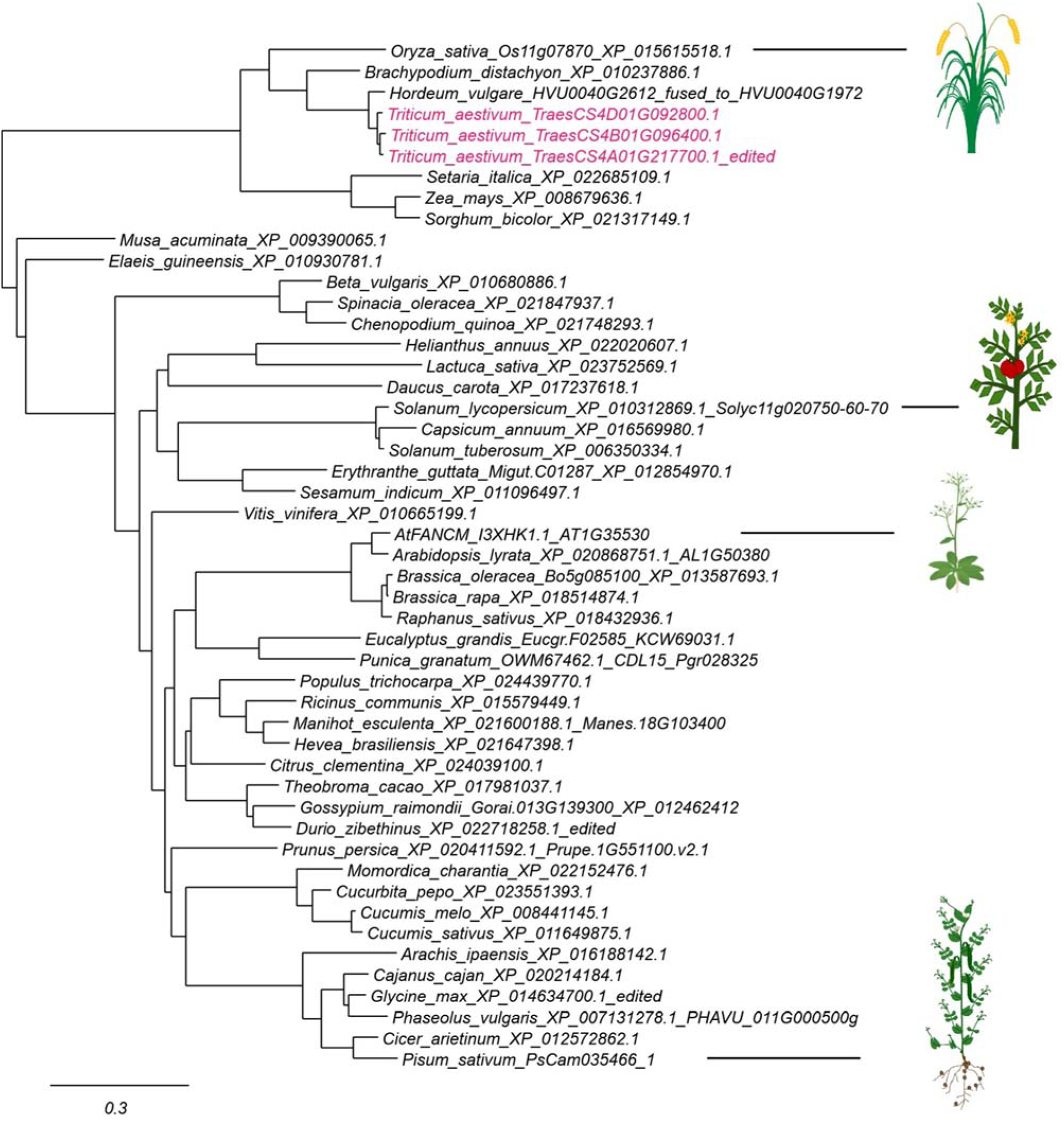
Phylogenetic tree of plant FANCM proteins. Genes present in several copies in a given species have been colored. Proteins sequences and accession numbers can be found in dataset S1

**Figure S3.**
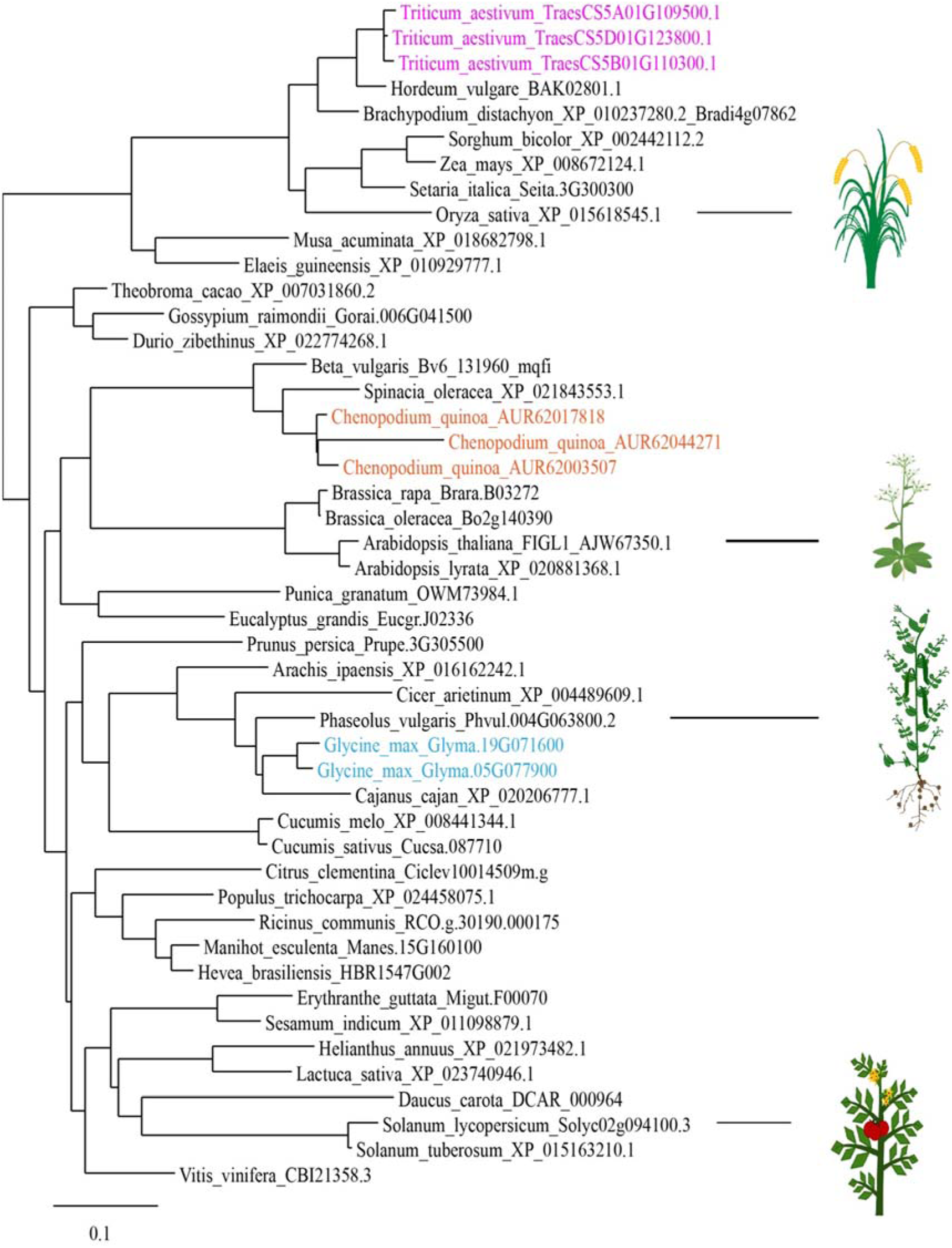
Phylogenetic tree of FIGL1. Genes present in several copies in a given species have been colored. Proteins sequences and accession numbers can be found in dataset S1

**Figure S4:**
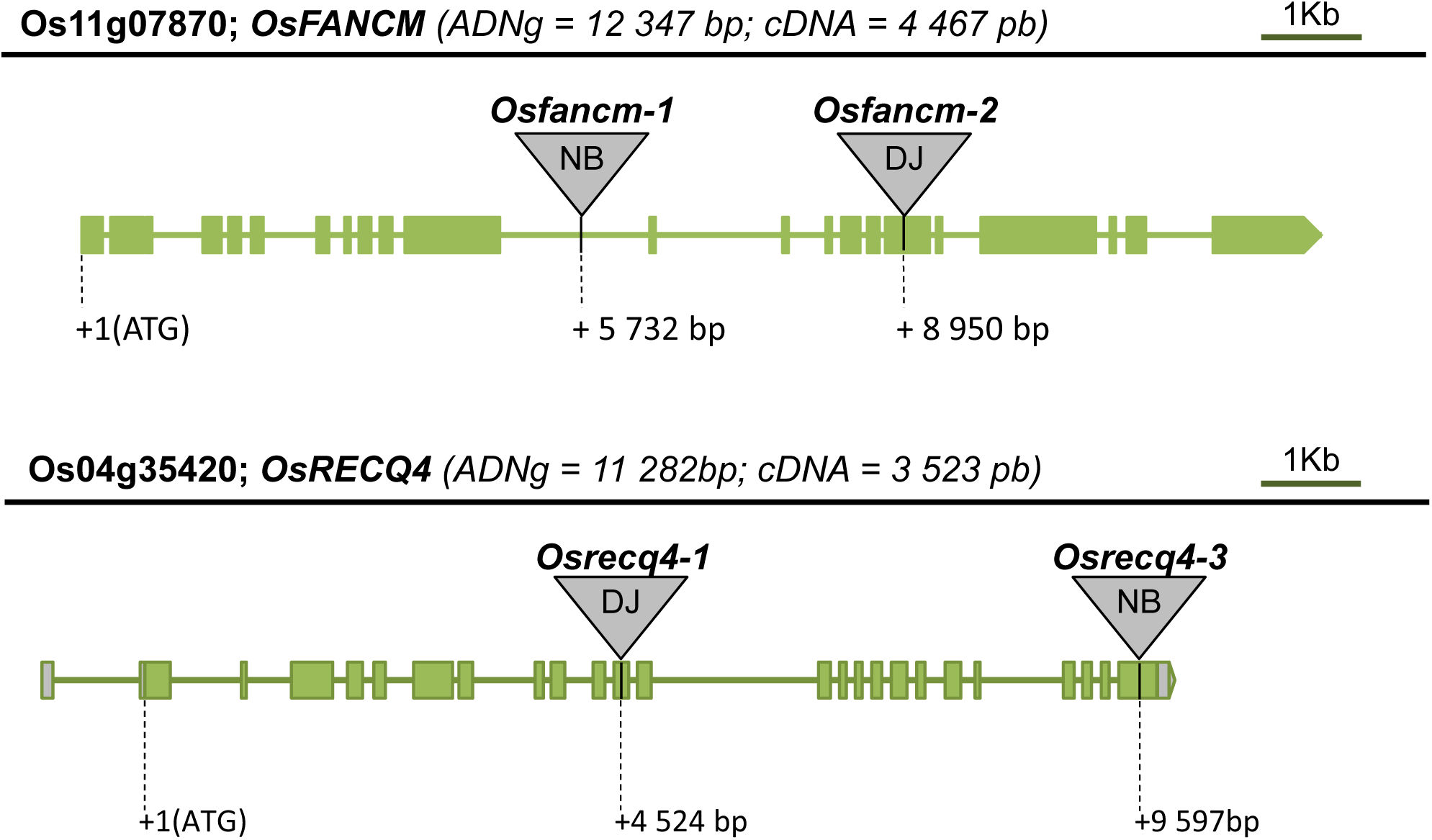
Positions of T-DNA insertions in *OsFANCM* and *OsRECQ4*. T-DNA insertions are indicated with a triangle. Mutants are from two different cultivars, Nipponbare (NB) or Dongjin (DJ). The exact position of the T-DNA insertion site was confirmed by Sanger sequencing.

**Figure S5.**
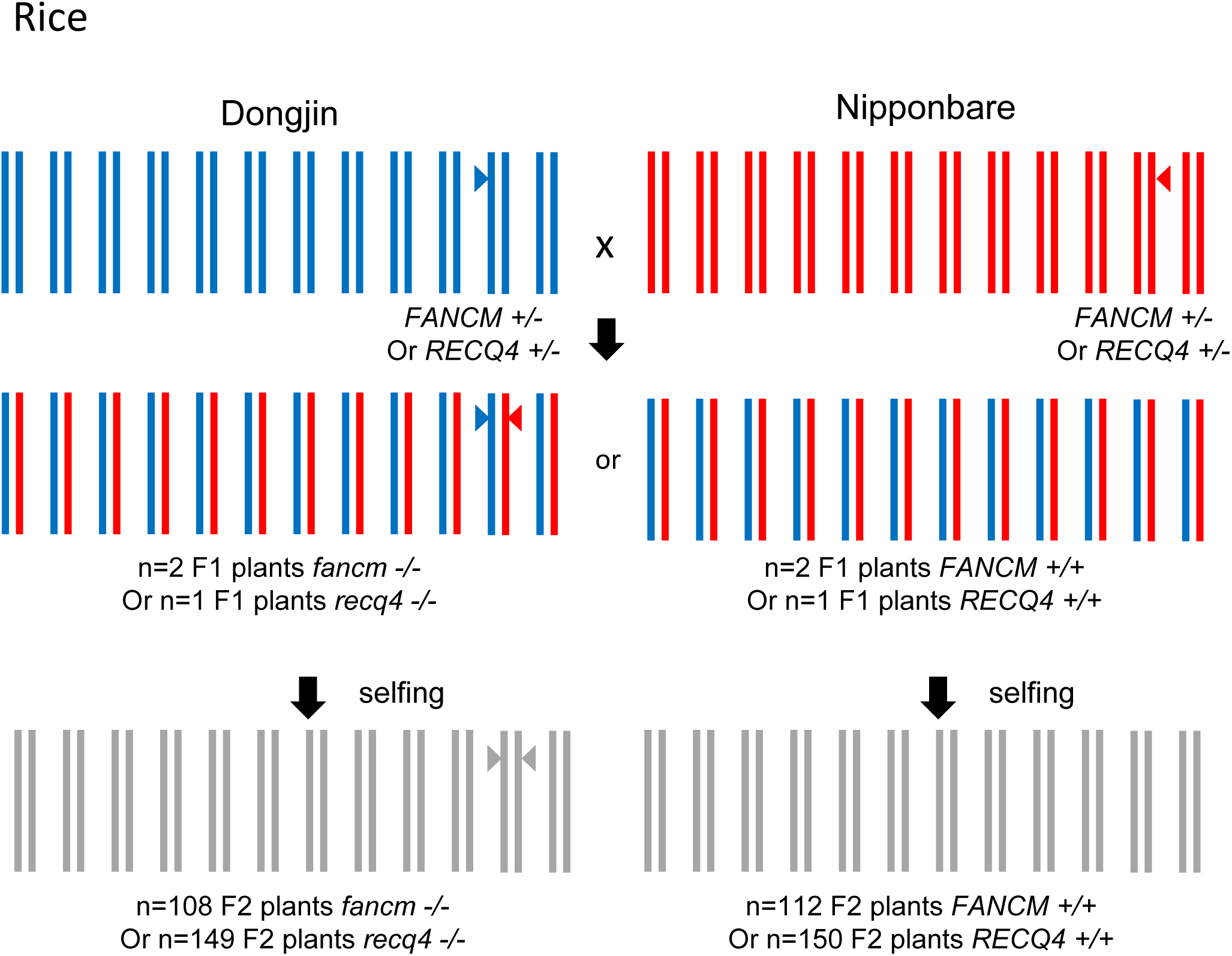
Experimental scheme for rice *fancm* or *recq4* Dongjin/Nipponbare hybrid populations.

**Figure S6.**
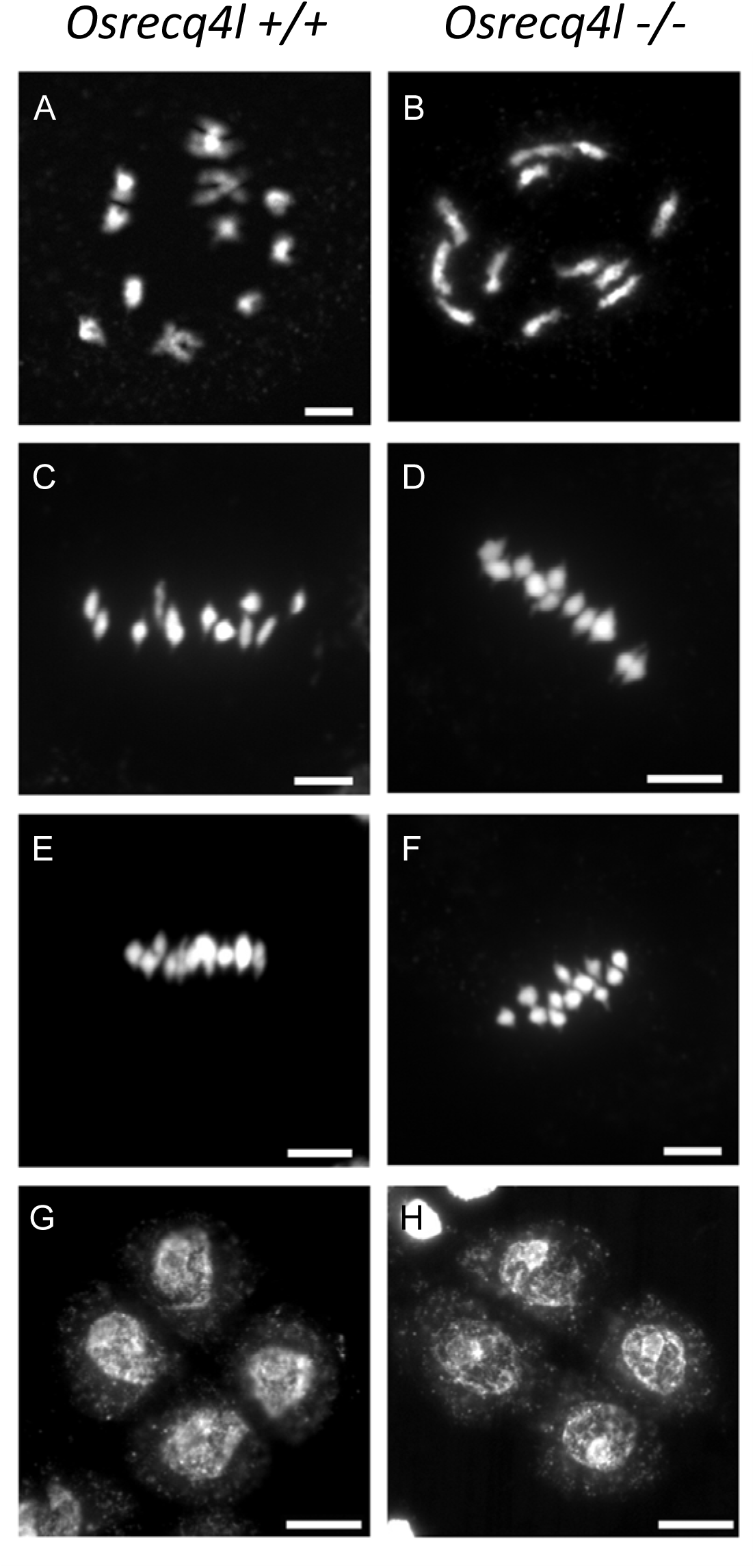
Male meiosis in *Osrecql4 -/-* (A-B) Diplotene, the 12 pairs of chromosome are connected by chiasma. (C-D) Metaphase I with 12 aligned bivalents. (E-F) Metaphase II with 12 pairs of chromatids. (G-H) Telophase II. Male meiotic chromosome spreads were performed as previously described in [17]. Scale bar = 5 μm.

**Figure S7.**
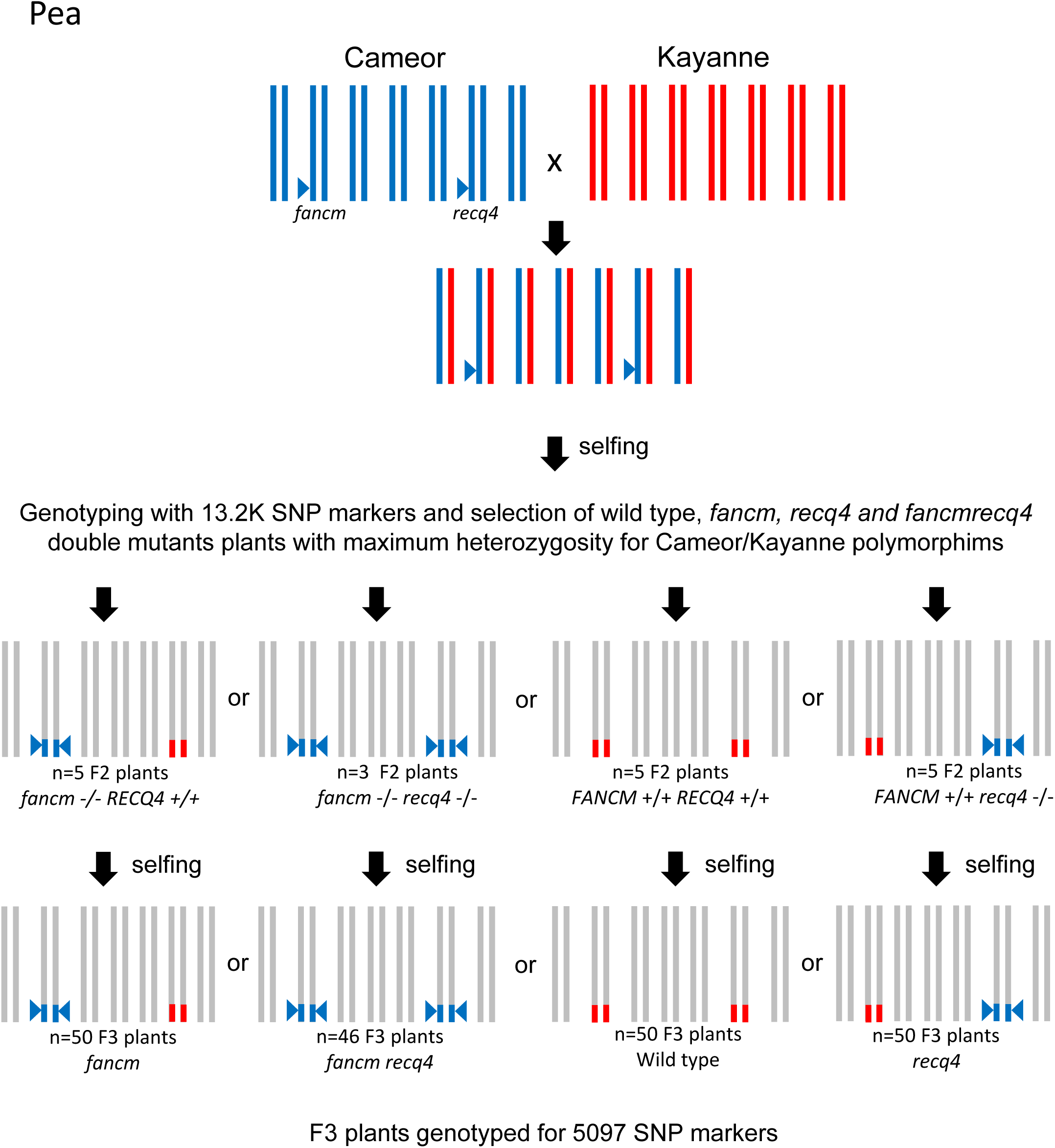
Experimental scheme for Pea *recq4 fancm* Cameor/Kayanne hybrid population

**Figure S8.**
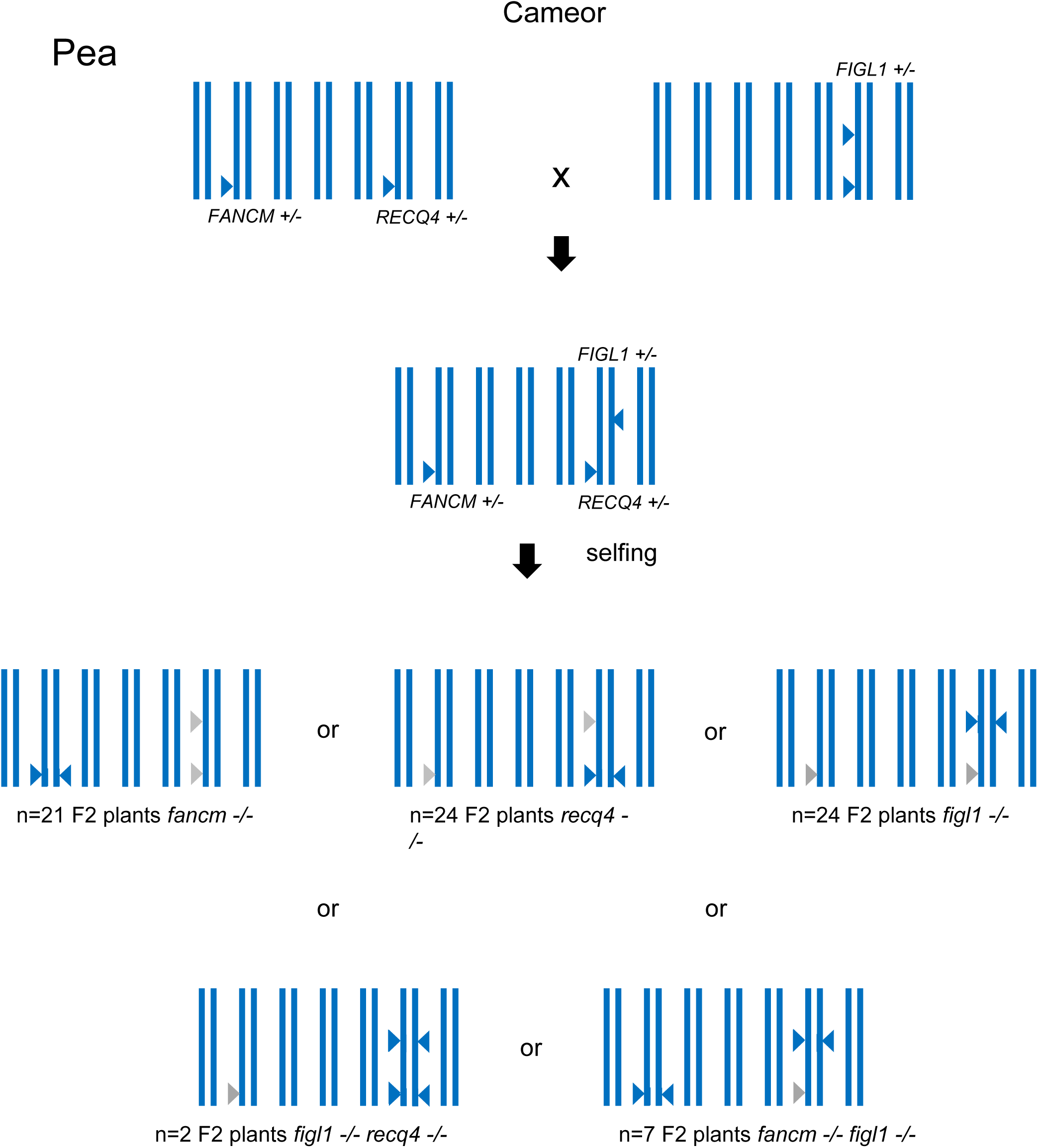
Crossing scheme for the Pea *recq4 fancm figl1* Cameor population.

**Figure S9.**
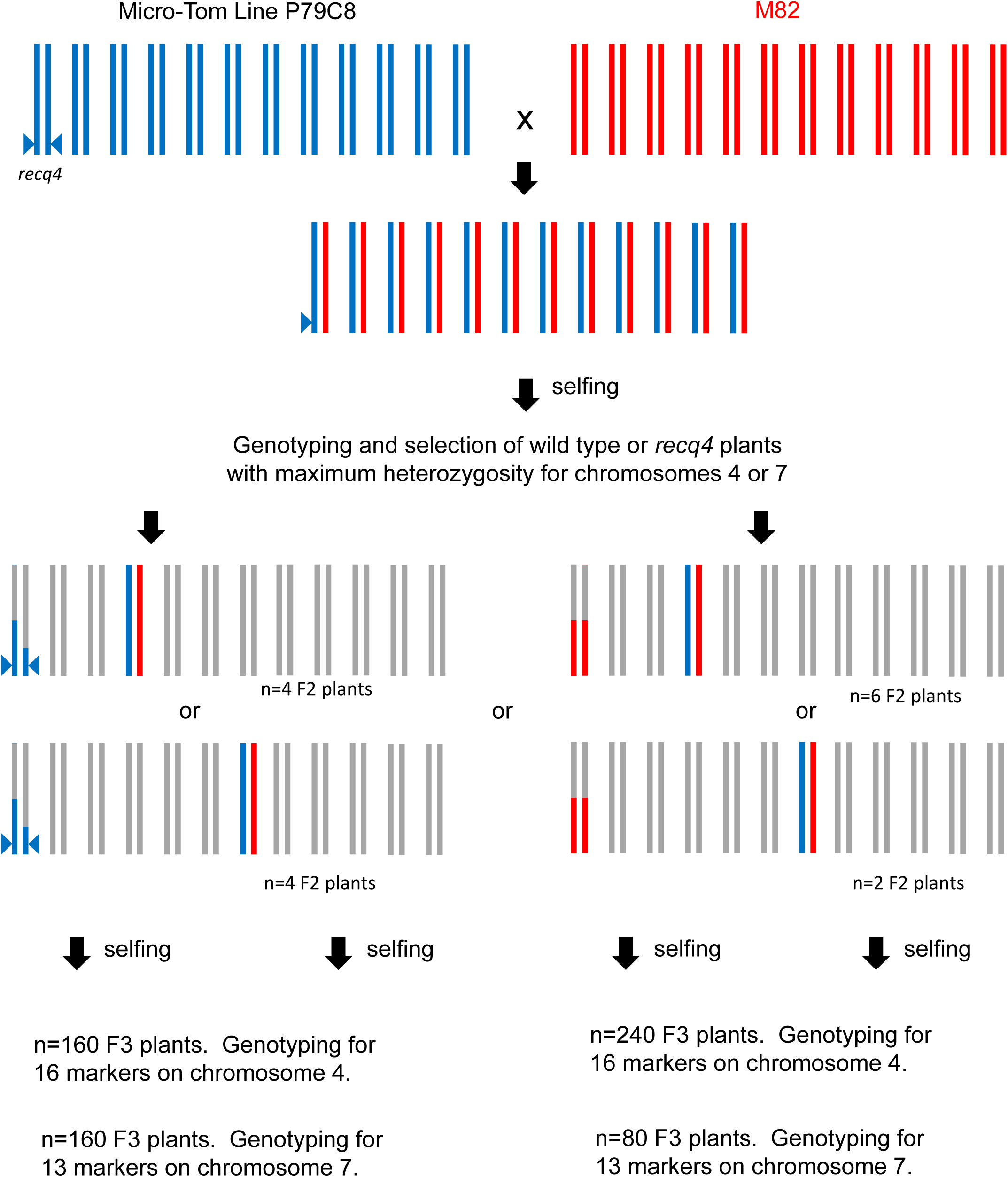
Experimental scheme for Tomato *recq4* Micro-Tome/M82 hybrid population

**Table S1:**
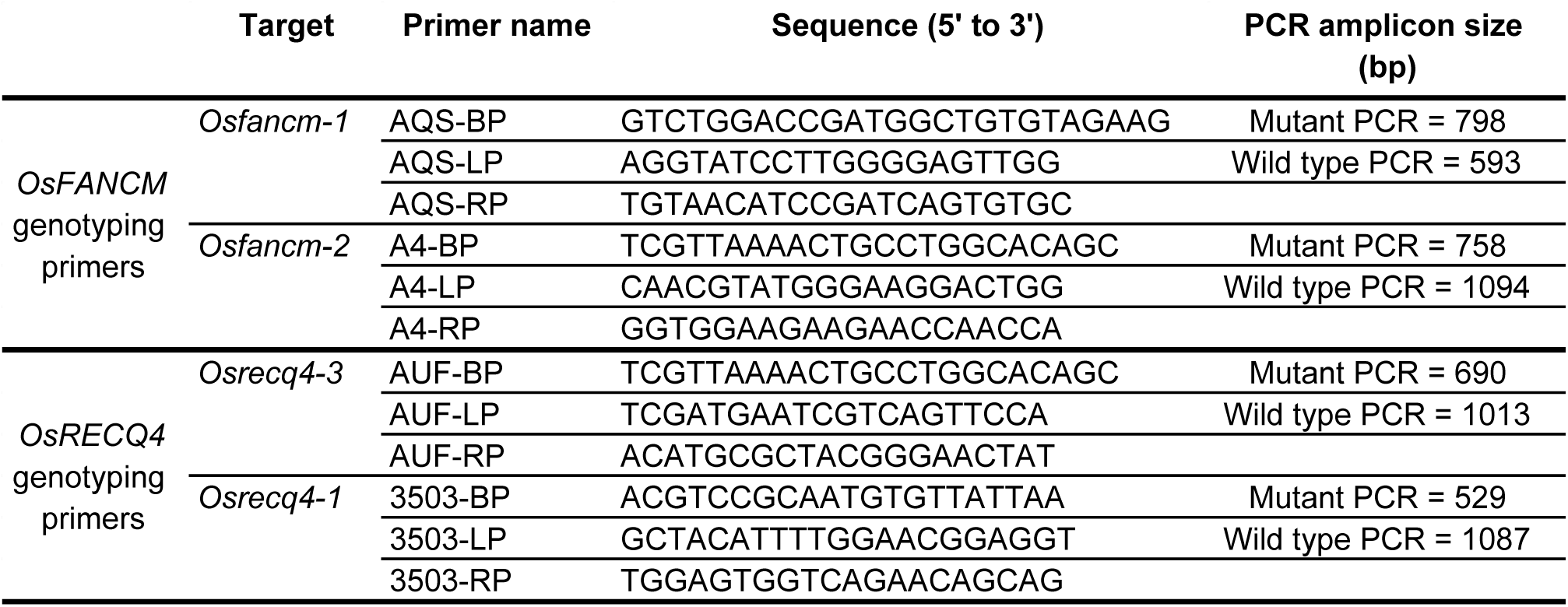
Primer sequences used for genotyping rice mutants and SNP position in the rice genome (MSU v7.0). Wild type PCR was done with LP (Left primer) and RP (Right primer) primers; Mutant PCR with BP (Backbone primer) and RP primers.

